# Environmental Gradients Shape the Hydrocarbon-Degrading Microbiome in Two Mid Atlantic Bays

**DOI:** 10.64898/2026.03.25.714183

**Authors:** Dinuka L. J. Patabandige, Jojy John, Maximiliano Ortiz, Barbara J. Campbell

**Affiliations:** Department of Biological Sciences, Clemson University, Clemson, SC, USA; Clemson University Genomics & Bioinformatics Facility, Clemson University, Clemson, SC, USA

**Keywords:** Microbial hydrocarbon degradation, Metagenome, Metatranscriptome, metagenome assembled genome (MAGs)

## Abstract

Hydrocarbons are recalcitrant organic matter that are released into the environment via natural and anthropogenic activities. We hypothesized that abiotic and biotic factors, including salinity, temperature, seasonality, microbial interactions, and functional redundancy, influence the abundance and activity of potential hydrocarbon degraders in the Delaware and Chesapeake Bays. We identified key genes in hydrocarbon degradation pathways in metagenomes, metatranscriptomes, and metagenome assembled genomes (MAGs) from these estuaries. Aerobic aromatic and alkane degradation pathways predominated in both estuaries, with higher gene abundances observed in low-salinity spring and summer samples. Hydrocarbon degrading MAG abundance were significantly structured by salinity, temperature, nitrate, and silicate concentrations. Metatranscriptomic analyses revealed consistently higher expression of aerobic alkane and aromatic degradation genes in the Delaware compared to the Chesapeake Bay, with the highest occurring under low-salinity spring conditions in the former. Catechol degradation pathways exhibited high functional redundancy, whereas the naphthalene degradation pathway showed restricted distribution. Co-expression analysis revealed that *Burkholderiales* displayed condition dependent metabolic coupling while *Pseudomonadales* integrated hydrocarbon degradation with fermentation and central metabolism, demonstrating complementary strategies that support multi-scale ecosystem resilience. In conclusion, environmental gradients and taxon-specific metabolic strategies together govern hydrocarbon degradation potential in these estuaries, with implications for predicting ecosystem responses to hydrocarbon inputs under changing conditions.

**Importance:** Coastal estuaries are among the most contaminated aquatic environments on Earth, receiving continuous hydrocarbon inputs from industrial activity, urban runoff, and natural sources. Microorganisms are the primary agents of hydrocarbon breakdown in these systems yet predicting when and where this capacity is active and how resilient it is to environmental change remains a major challenge. Using paired genomic and transcriptomic data from microbial genomes across two major mid-Atlantic estuaries, we show that hydrocarbon degradation capacity is not uniformly distributed but is instead shaped by salinity, nutrients, and seasonality in pathway-specific ways. Critically, dominant degrader taxa employ fundamentally different metabolic strategies to sustain this function across fluctuating conditions, providing a form of community-level insurance against environmental disturbance. These findings advance our ability to predict microbial hydrocarbon degradation in coastal systems and inform nature-based approaches to bioremediation under increasing climate and anthropogenic pressures.

## Introduction

Estuarine ecosystems represent dynamic interfaces between terrestrial and marine environments, where complex biogeochemical processes govern the fate of organic matter. These transitional zones sustain high biological productivity and provide crucial ecosystem services, yet they are affected by the accumulation of hydrocarbons, including polycyclic aromatic hydrocarbons (PAHs), from both natural and anthropogenic sources (1). Natural inputs such as terrestrial plant material, diagenetic processes, and hydrocarbon seeps contribute to background PAH levels (2), while oil spills, industrial discharges, and urban runoff introduce additional loads (1, 3). These compounds can persist in sediment and water, posing ecological risks (4) Understanding the fate of these contaminants is essential for protecting estuarine biodiversity, maintaining ecosystem functions, and mitigating long-term environmental impacts (5, 6).

Microbes play a pivotal role in hydrocarbon degradation, and their activity and composition are highly sensitive to environmental gradients such as salinity, redox potential, and nutrient availability (1, 7). Many diverse Bacteria and Archaea can metabolize hydrocarbons through aerobic and anaerobic pathways, transforming these pollutants into less harmful compounds (7). About 79 bacterial genera within the classes of Alphaproteobacteria, Gammaproteobacteria, Betaproteobacteria, Actinobacteria, and Bacteriodia have been identified to potentially degrade hydrocarbons (1, 8). Hydrocarbon-degrading bacteria employ diverse catabolic pathways, including alkane hydroxylation (9), aromatic ring activation via mono- and dioxygenases, and ring-cleavage reactions mediated by ring-hydroxylating dioxygenases (10). Together, these pathways facilitate the transformation of aliphatic (9), monoaromatic (11–13), and polycyclic aromatic hydrocarbons (1, 14) into central metabolic intermediates that can be assimilated for growth.

The bioavailability of hydrocarbons is limited by hydrophobicity, low solubility and sorption of suspended particles, mineral or organic material, especially in the case of higher molecular weight PAHs (1, 15, 16). The rate of reaction is affected by chemical complexity but may be increased by functional redundancy or microbial co-metabolism (17). Temperature affects both hydrocarbon properties and enzymatic activity of microbial degradation, with biodegradation generally occurring at temperatures of 15-30 °C (16, 18) and drives seasonal changes in the diversity and functional potential of PAH degraders (19). Nutrient availability, in particular carbon, nitrogen and phosphorus balance, controls degradation; nutrient scarcity may inhibit activity due to competition with primary producers (17, 20–22), while nutrient surplus may inhibit microbial growth (5). The availability of oxygen is a key factor, since aerobic degradation pathways are generally faster (14, 23), while anaerobic processes relying on alternative electron acceptors are slower because of lower energy yields (11, 23, 24). Salinity variations affect microbial activity and community composition, low salinity favors greater diversity and higher numbers of the hydrocarbon degradation genes (25, 26), while high salinity favors tolerant taxa such as *Acinetobacter* and *Halomonas,* which can be dominant but often with low degradation efficiency (8, 27).

The Chesapeake and Delaware Bays are estuarine systems in the mid-Atlantic U.S. that exhibit distinct hydrographic and biogeochemical characteristics that shape microbial community structure and hydrocarbon degradation processes (28). The Chesapeake Bay is partially mixed and characterized by substantial freshwater input, particle deposition, and seasonal hypoxia in deeper waters driven by phytoplankton blooms, nutrient influx, and salinity stratification (26, 29). Critically, these phytoplankton blooms not only drive hypoxic conditions but also contribute biogenic aromatic compounds, including PAHs, directly linking primary production dynamics to hydrocarbon cycling in this system (30, 31). The resulting redox stratification promotes a shift from aerobic to slower anaerobic degradation pathways, potentially facilitating PAH accumulation in deeper, oxygen-depleted waters. In contrast, Delaware Bay exhibits a well-mixed water column with a steep salinity gradient, high turbidity, and limited primary production despite significant nutrient inputs (32, 33), reducing biogenic PAH contributions while maintaining more consistent aerobic degradation conditions. Both bays nevertheless accumulate aromatic compounds from anthropogenic inputs in addition to natural sources such as salt marsh vegetation and phytoplankton (28, 30, 32), with surface waters of Chesapeake Bay showing elevated PAH levels linked to human activity, though clear seasonal patterns are lacking (34). *Alphaproteobacteria, Betaproteobacteria, Gammaproteobacteria, Acidimicrobiia,* and *Actinomycetia* are prominent hydrocarbon degraders in both bays (21, 35–38). Functional redundancy (FRed), the ability of phylogenetically different taxa to perform equivalent metabolic functions, can maintain hydrocarbon turnover in both systems despite different environmental conditions (39). This overlap in catabolic potential between degradation guilds may separate the composition of the community from the function of the ecosystem, potentially masking significant differences in degradation efficiency between the different bays (40, 41).

Despite knowledge of microbial taxa capable of hydrocarbon degradation, significant gaps remain in understanding how spatial and temporal environmental variability shapes the composition, functional potential, functional redundancy and gene expression of these communities in estuarine surface waters, particularly in relation to hydrocarbon transformation pathways. In this study, we integrated metagenomics, metatranscriptomics, and MAG-resolved co-expression analysis across metagenomes and metatranscriptomes spanning two contrasting estuaries, multiple seasons, and a broad salinity gradient to examine how environmental gradients and taxon-specific metabolic strategies jointly govern hydrocarbon degradation potential at the genome level. We investigated the extent to which salinity, nutrients, and seasonality structure the distribution and transcriptional activity of hydrocarbon-degrading communities, whether functional redundancy encoded in metagenomic potential is realized through simultaneous active expression across multiple lineages in situ, and how dominant degrader taxa differs in their metabolic integration of hydrocarbon catabolism with broader cellular processes. Together, these analyses provide a genome-resolved transcriptional framework for understanding and predicting how estuarine microbial communities sustain hydrocarbon degradation capacity under shifting environmental conditions.

## Results

### Abundance and Functional Potential of Hydrocarbon-Degrading Bacteria

Having established the broad distribution of hydrocarbon degradation genes at the community gene level from assembled contigs (Supplementary information, Figure S1), we sought to identify the taxonomic composition and genomic potential of organisms harboring these capabilities by examining 360 previously described bacterial MAGs (>70% complete, < 5% contamination) from the Delaware and Chesapeake Bays, dereplicated to the Order level (38, 41). After screening, 181 MAGs encoded genes associated with hydrocarbon degradation pathways and comprised between 3-37% of all recovered MAGs (Figures 1, S2, Table S2). The relative abundance of hydrocarbon-related MAGs varied across seasons, bays, and salinity classes. Spring samples showed the highest overall MAG relative abundances, with values reaching up to ∼37% in mid-salinity Delaware Bay samples, while Chesapeake Bay samples showed comparatively lower abundances. Summer and fall samples displayed lower abundances relative to spring, though mid and high-salinity Delaware Bay samples maintained comparatively higher MAG abundances across these seasons. Community composition shifted across environmental gradients, with *Burkholderiales* and *Acidimicrobiales* associated with low to mid-salinity conditions, while *Pseudomonadales*, *Rhodobacterales*, and *Flavobacteriales* were more prevalent under high-salinity conditions. These MAGs represented 41 taxonomic orders and collectively contained 44 out of 116 key genes identified by KEGG (42) linked to hydrocarbon degradation (Figure 3). The genes collectively spanned aliphatic and aromatic hydrocarbon pathways and encompassed both aerobic and anaerobic processes.

**Figure 1.**
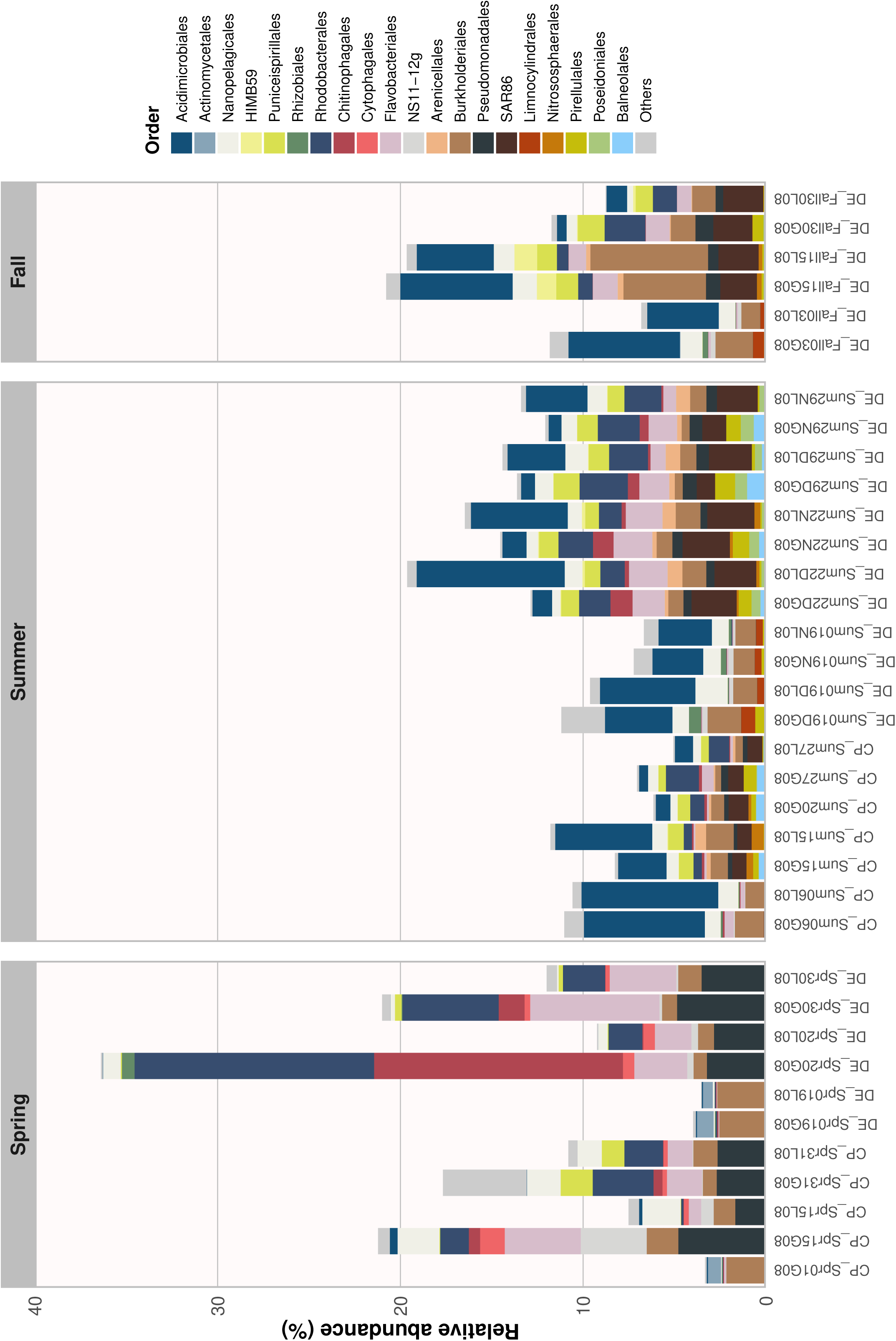
Relative abundance of top 20 MAGs across all samples based on CPM normalized counts, shown as stacked bar charts scaled 0 % to 40% grouped by season. *Abbreviations:* CP = Chesapeake Bay; DE = Delaware Bay; Spr, Sum, Fall = Spring, Summer, Fall; G08 = >0.8 μm; L08 = <0.8 μm; CPM= counts per million. Abundance of 181 MAGs and the phylogenomic tree are provided in Supplementary Figure 1 and Table 1.

At the broad taxonomic scale, the most prominent orders containing hydrocarbon degrading genes included *Burkholderiales, Acidimicrobiales, Flavobacteriales, Nanopelagicales, Rhodobacterales,* and *Pseudomonadales,* each contributing > 14 MAGs with pathways for polycyclic aromatic hydrocarbon (PAH) degradation under both aerobic and anaerobic conditions (Figure 2, Table S3). Benzene and benzoate degradation genes were widely distributed among *Burkholderiales*, *Rhodobacterales*, *Pseudomonadales*, and *Actinomycetales.* Catechol ortho- and meta-cleavage pathway genes were particularly abundant and detected across multiple bacterial orders, including *Burkholderiales*, *Pseudomonadales*, *Rhodospirillales*, *Actinomycetales*, and *Flavobacteriales*. Cumate and cymene degradation pathways were comparatively restricted, occurring mainly in *Actinomycetales* and *Rhodobacterales*, while phthalate degradation genes were most frequent in *Burkholderiales* and *Rhodobacterales*. Naphthalene degradation genes were confined to *Burkholderiales*. Both aerobic variants of toluene and xylene degradation genes were primarily detected in *Burkholderiales*, *and Acidimicrobiales*. Genes associated with anaerobic toluene were detected only sporadically, mainly within *Acidimicrobiales*.

**Figure 2.**
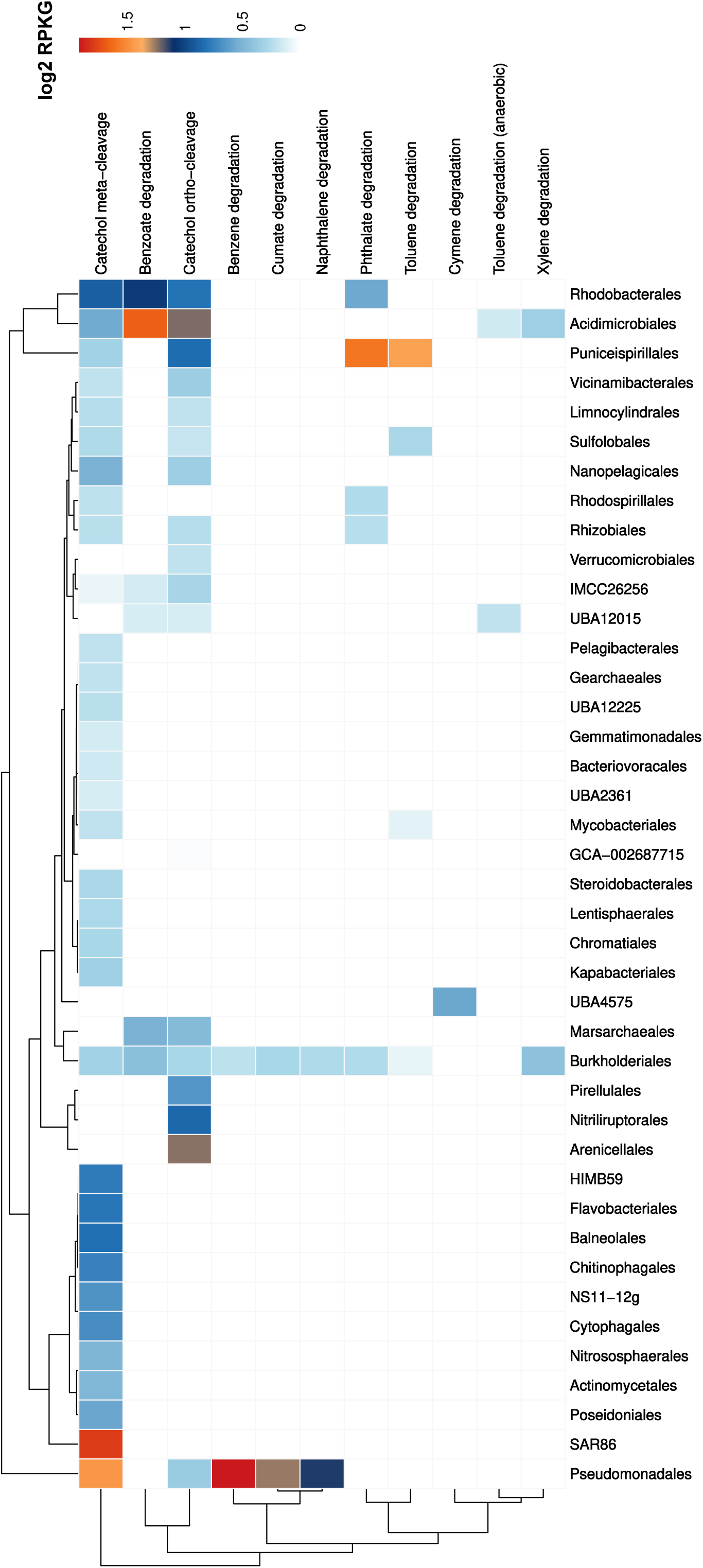
Mean abundance of hydrocarbon degradation pathways across metagenome-assembled genomes (MAGs) at the Order level. Quantified as log_2_RPKG values, which were calculated per gene, summed across all genes assigned to each pathway per MAG per sample.

Potential FRed varied substantially among aromatic degradation pathways (Table S3). Catechol degradation pathways exhibited the highest potential redundancy, with meta-cleavage pathway genes detected across 33 taxonomic orders (121 MAGs) and ortho-cleavage genes across 18 orders (75 MAGs). Benzoate and phthalate degradation genes showed high potential redundancy (6 orders/11 MAGs and 5 orders/16 MAGs, respectively), while toluene degradation displayed moderate redundancy (4 orders, 4 MAGs). In contrast, several pathways showed restricted potential such as xylene, naphthalene, benzene, cumate, and anaerobic toluene degradation were each limited to 2 taxonomic orders (3, 6, 3, 8, and 5 MAGs, respectively). Cymene degradation exhibited no potential redundancy, with genes confined to a single MAG.

Within all MAGs, hydrocarbon degradation genes exhibited pronounced variability in abundance across sites, seasons, and salinity conditions, reflecting shifts in microbial functional potential (Figure 3). Aerobic aromatic degradation genes were the most dominant group overall, with *pcaD* showing the highest abundance, reaching its peak in summer low salinity samples. Similarly, *dmpC/xylG/praB* and *mhpE* were most abundant in Delaware spring medium and Chesapeake summer low salinity samples, respectively, while *dmpB/xylE* and genes involved in phthalate degradation, such as *pht5*, were also highly represented, with the greatest abundances consistently observed in Delaware summer medium salinity samples.

**Figure 3.**
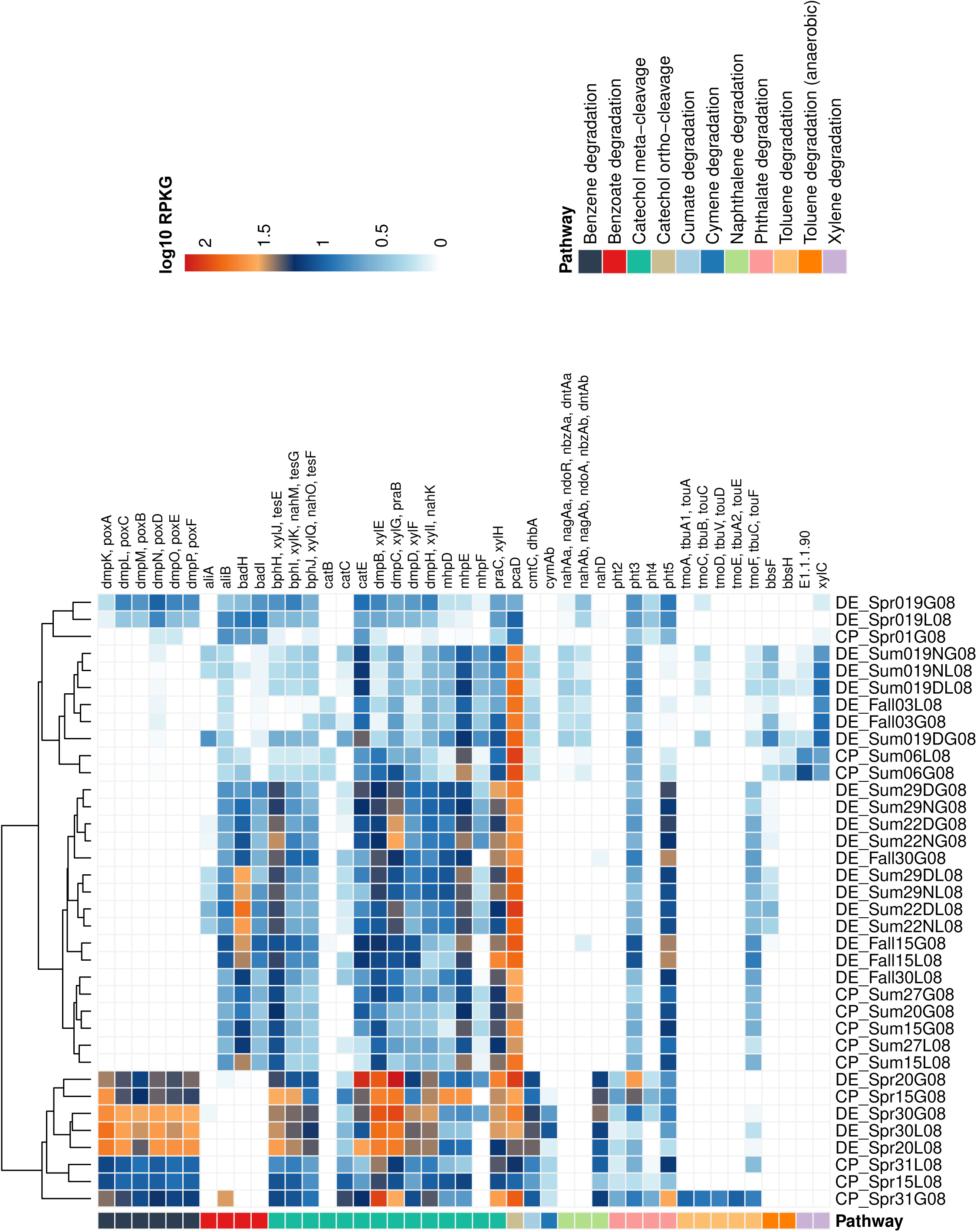
Abundance of hydrocarbon degradation genes within MAGs across the samples, quantified as log_10_ RPKG. *Abbreviations:* CP = Chesapeake Bay; DE = Delaware Bay; Spr, Sum, Fall = Spring, Summer, Fall; G08 = >0.8 μm; L08 = <0.8 μm; RPKG = reads per kilobase per genome equivalent. Full gene names, enzyme functions, and pathway details are provided in Supplementary Table 2.

### Environmental Drivers Shape the Distribution and Abundance of Hydrocarbon Degrading Bacteria

A distance-based redundancy analysis (db-RDA) based on Bray–Curtis dissimilarities revealed that environmental parameters collectively exerted a strong influence on the hydrocarbon gene encoding MAG community compositions across the estuarine gradient (Figure 4, Table S4). The overall model was significant (F = 10.24, *p* = 0.001), indicating that the measured physicochemical variables explained a substantial proportion of the observed compositional variation. Among individual factors, nitrate, temperature, salinity, and silicate emerged as the strongest drivers of MAG distribution patterns, while phosphate, bacterial production, and cell abundance also contributed significantly. The first two canonical axes were strongly significant (both *p* = 0.001) and captured the dominant environmental gradients structuring microbial communities, especially salinity and temperature. Taxonomic associations, inferred from ordination positions, reflected these environmental gradients. *Rhodobacterales*, *Pseudomonadales*, and *Chitinophagales* MAGs were positively associated with high-salinity conditions. In contrast, *Acidimicrobiales* MAGs were aligned with lower salinity and elevated temperature. *Actinomycetales* and *Nanopelagicales* MAGs were positioned along a low-salinity and higher nitrate and phosphate concentrations.

**Figure 4.**
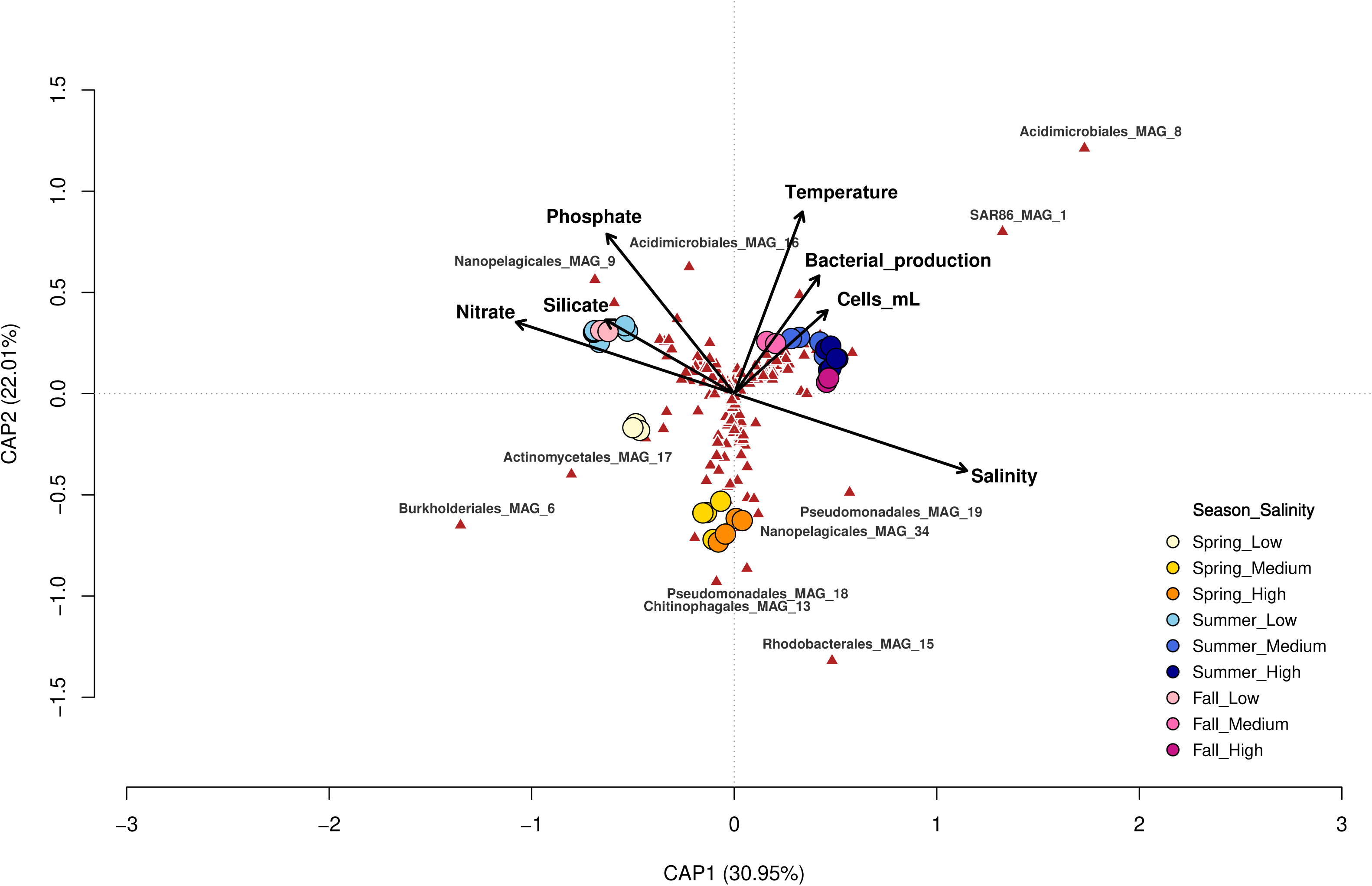
Distance-based redundancy analysis of hydrocarbon-degrading MAGs from Chesapeake and Delaware Bay samples. Points represent samples grouped by season (Spring, Summer, Fall) and salinity category (Low, Medium, High). Vectors indicate significant environmental predictors, including salinity, bacterial production, cell density, temperature, nitrate, phosphate, and silicate. MAGs with the strongest associations with environmental axes are shown as biplot loadings.

To assess whether these same gradients shape hydrocarbon degradation potential, we examined correlations between environmental variables and degradation genes from MAGs grouped by functional pathway (Figure 5, Table S2). In the catechol meta-cleavage pathway, genes including *bphH, bphI, dmpB, dmpC, dmpD, mhpD*, and *praC,* exhibited strong positive correlations with salinity and strong negative correlations with temperature, silicate, nitrate, and phosphate concentrations. However, *bphJ* was negatively associated with temperature, while *mhpE* was positively associated with temperature and negatively associated with ammonium. Catechol ortho-cleavage genes displayed significant positive correlations with phosphate and negative correlations with silicate concentrations. In the benzoate and benzene degradation pathways, *badH and dmpN* exhibited positive correlations with salinity and phosphate concentrations, while *dmpN* was negatively correlated with salinity. Temperature showed strong negative correlations with *dmpK, dmpL, dmpM,* and *dmpO*. Nitrate and phosphate concentrations exhibited strong positive correlations while salinity exhibited strong negative correlations with genes in the naphthalene degradation pathway *(nahAb, nagAb, ndoA, nbzAb,* and *dntAb*). For xylene degradation, salinity was negatively correlated with *xylC* and positively correlated with nitrate and phosphate concentrations. The anaerobic toluene degradation genes *bbsF* and *bbsH* showed strong positive correlations with phosphate concentrations and negative correlations with salinity. In contrast, the aerobic toluene degradation gene *tmoC* exhibited a strong negative correlation with salinity and positive correlations with nitrate and phosphate concentrations, while *tmoF* showed a positive correlation with salinity and a negative correlation with nitrate concentrations. Phthalate degradation genes showed significant negative correlations with multiple variables: temperature, phosphate and silicate concentrations (*pht2*), and temperature (*pht4*). However, *pht5* showed a strong positive correlation with salinity and negative correlations with nitrate and phosphate concentrations.

**Figure 5.**
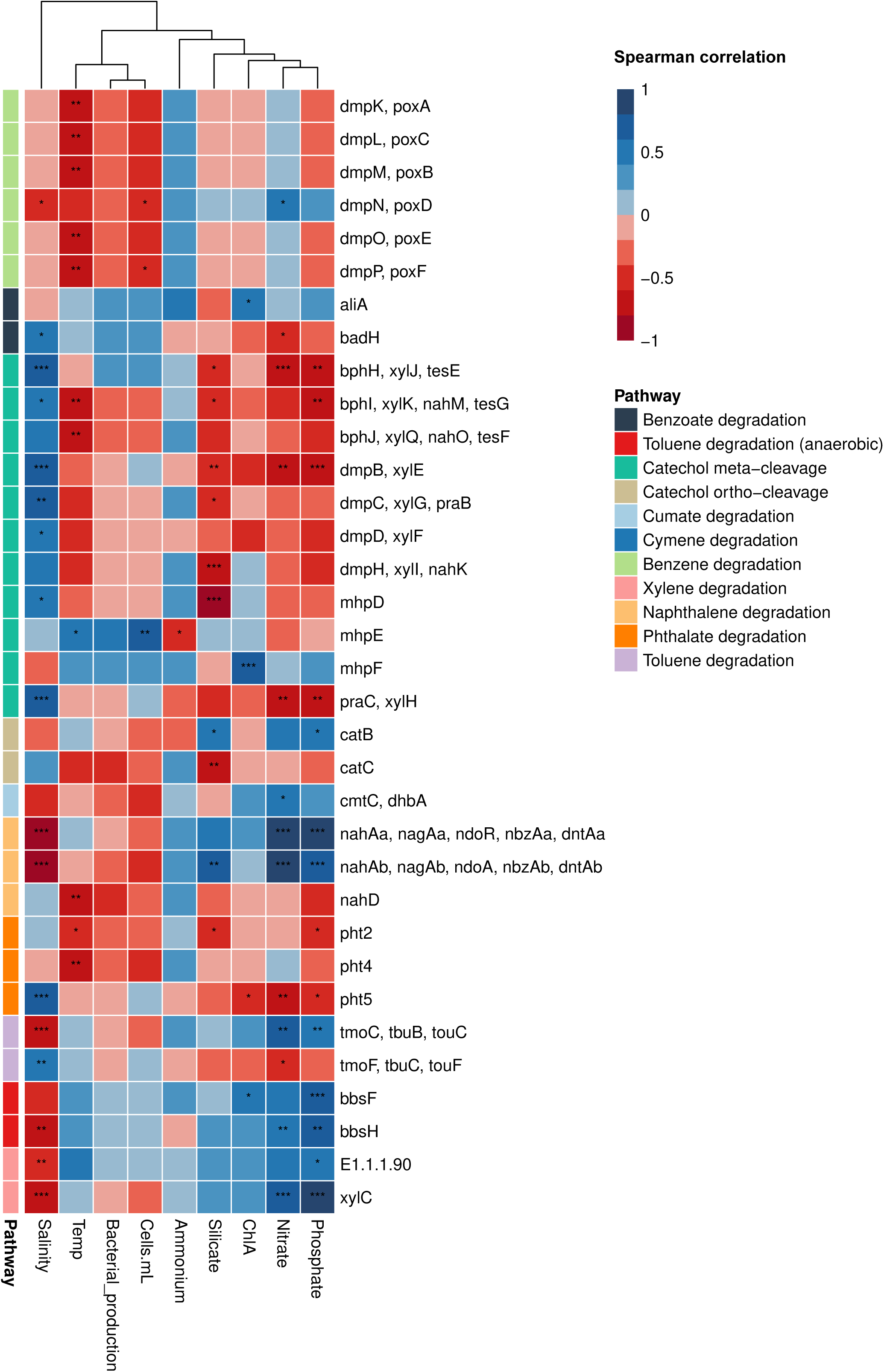
Correlation matrix showing associations between abundance of hydrocarbon degradation genes and environmental metadata variables across all samples. Spearman correlation coefficients indicated and asterisks denote significance levels (*p* < 0.05 *, p* < 0.01 \**, p* < 0.001 ***). Genes (detailed in Supplementary Table 2) are grouped by metabolic pathway. *Abbreviation*: chlorophyll *a* (ChlA), Temperature (Temp).

### *In Situ* Activity and Redundancy of Bacterial Hydrocarbon Degraders

Consistent with community level metatranscriptomic patterns (Supplementary Information, Figure S3), genome resolved expression analysis revealed distinct hydrocarbon degradation gene activity across bay type, salinity gradients, and seasons (Figure 6). Gene expression patterns were clearly separated between bays, with Delaware Bay samples exhibiting higher expression across most hydrocarbon degradation pathways. Within Delaware Bay, both season and salinity influenced gene expression. Seasonally, spring samples had the highest expression of catechol and benzene degradation genes (Figure 6). Summer samples maintained elevated catechol gene expression, while fall samples retained moderate benzoate degradation expression alongside sustained catechol gene activity. Salinity gradients further modulated expression, with low salinity samples exhibiting the highest relative expression of benzene and catechol degradation genes and progressive declines through medium to high salinity conditions. Within Chesapeake Bay, gene expression was generally lower but displayed distinct seasonal and salinity trends. Spring samples showed moderate to higher expression across benzene, catechol, and phthalate degradation genes, while summer samples exhibited reduced expression with moderate activity limited to benzoate, phthalate, and catechol genes. Salinity effects mirrored those in Delaware Bay, low salinity samples showed relatively higher expression than medium or high salinity samples, but absolute expression levels remained lower than Delaware Bay across all salinity ranges.

**Figure 6.**
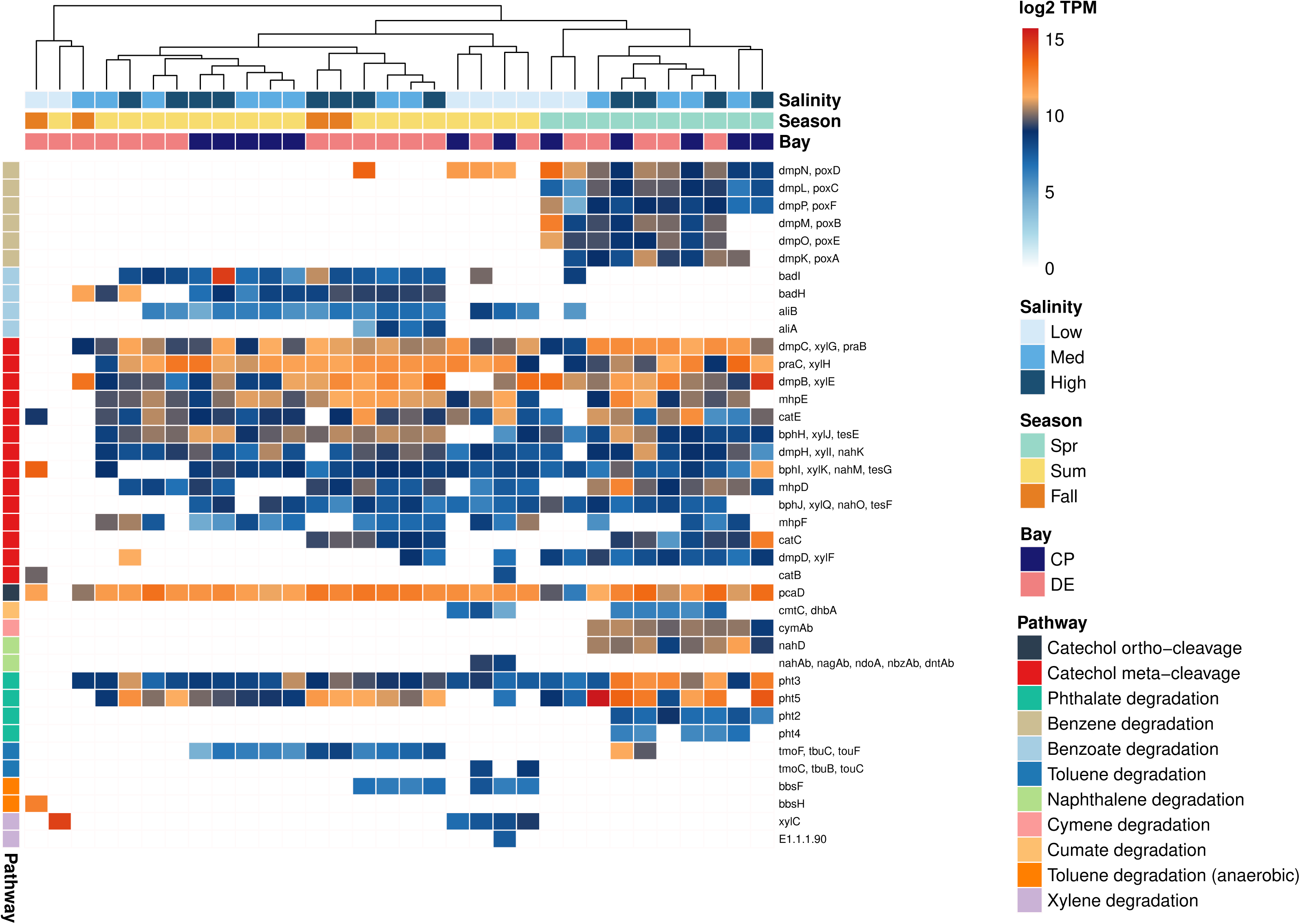
MAG-derived hydrocarbon degradation gene expression, quantified by log_2_ TPM-normalized transcript abundance values. *Abbreviations:* CP = Chesapeake Bay; DE = Delaware Bay; Spr, Sum, Fall = Spring, Summer, Fall; TPM = transcripts per million. Full gene names, enzyme functions, and pathway details are provided in Supplementary Table 2.

Realized FRed differed considerably across aromatic degradation pathways (Table S5). Meta-cleavage catechol degradation genes were most widely distributed, appearing in 121 MAGs spanning 29 orders, while ortho-cleavage genes occurred in 65 MAGs across 14 orders. Phthalate degradation demonstrated substantial redundancy (16 MAGs, 5 orders), whereas toluene and benzoate degradation showed intermediate distribution (4 and 3 orders, respectively). Conversely, naphthalene and anaerobic toluene degradation pathways exhibited narrow distribution, each restricted to 2 orders. Cymene degradation represented the most specialized pathway, with genes present from a single taxonomic order, indicating no realized FRed.

### Metabolic Potential and Co-expression Patterns in Hydrocarbon Degrading MAGs

Screening of hydrocarbon-degrading MAGs across six dominant orders for additional metabolic functions revealed heterogeneous metabolic potential (Figure 7a, Table S6). MAGs in all orders generally encoded key aerobic respiration genes (*atpA, coxA, cydA, nuoF*), except *Flavobacterales MAGs (*missing *nuoF) and Acidimicrobiales* and *Nanopelagicales* (missing *cydA*). Denitrification genes were detected sporadically in *Pseudomonadales*, *Flavobacteriales*, and *Rhodobacterales*, while sulfur metabolism genes were prevalent in *Flavobacteriales* and *Rhodobacterales*. Carbon fixation genes were largely absent, indicating reliance on heterotrophic metabolism, although most had rhodopsin related genes, which may be involved in phototrophy (43).

**Figure 7.**
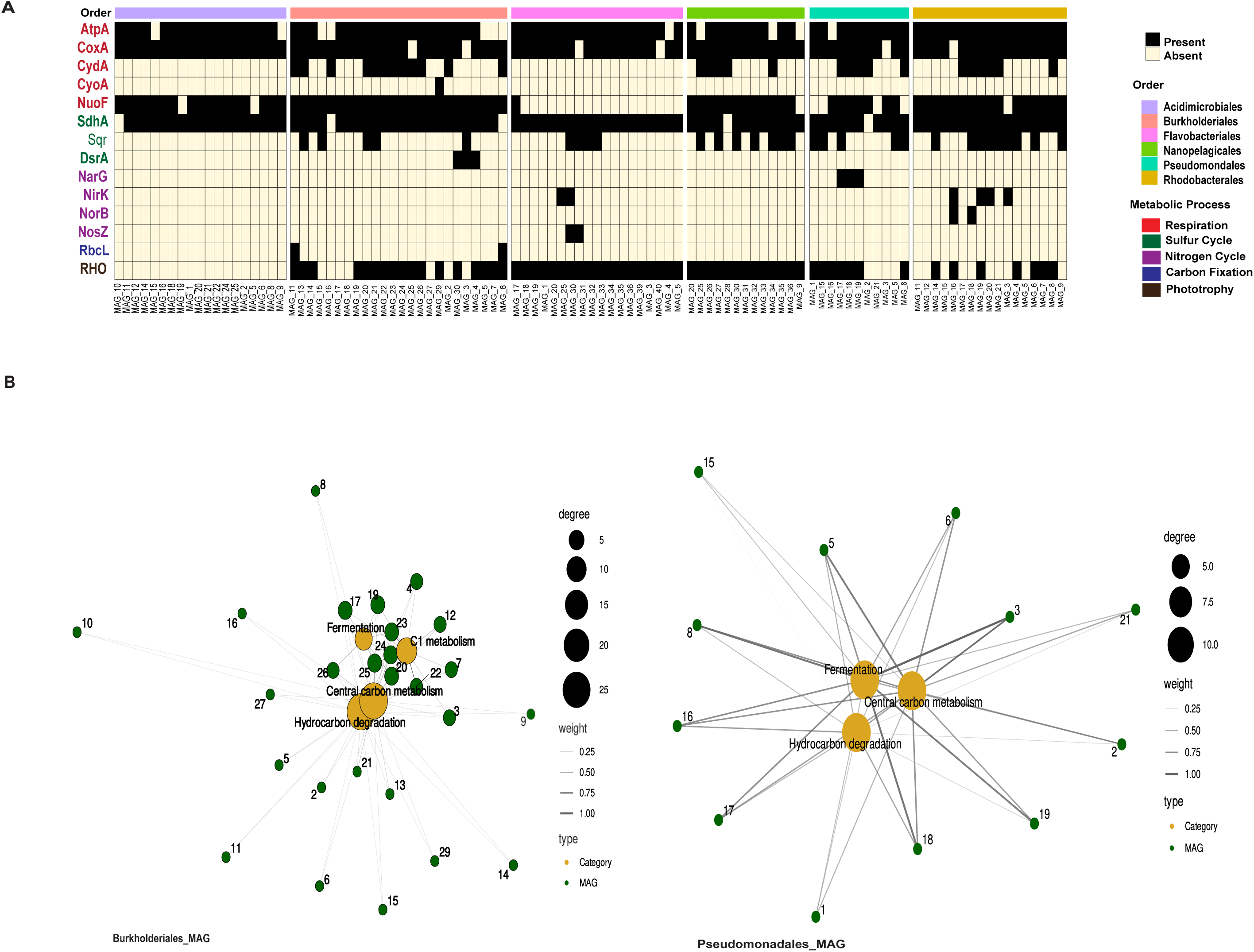
Metabolic potential and co-metabolic pathway associations in hydrocarbon-degrading MAGs. (a) Presence/absence heatmap of key metabolic marker genes across MAGs at the order level. (b) Co-occurrence networks for *Burkholderiales* (left, n = 26 MAGs) and *Pseudomonadales* (right, n = 12 MAGs). Gold nodes represent metabolic pathway categories and green nodes represent individual MAGs.

Network analysis (Figure 7b, Table S6) of metabolic pathway co-expression in *Burkholderiales* MAGs (n = 26) revealed a more heterogeneous network topology compared to *Pseudomonadales*, with MAGs varying in the number of connected metabolic categories (edges) ranging from two to four, where total edge weights ranged from 0.004 to 1.48 which suggesting greater variability in metabolic pathway integration across this order than in *Pseudomonadales*. Four metabolic categories were represented, C1 metabolism, central carbon metabolism, fermentation, and hydrocarbon degradation. C1 metabolism was the dominant category across the majority of MAGs (MAGs 3, 4, 5, 7, 12, 17, 19, 20, 22, 23, 24, and 25), while central carbon metabolism led in 13 MAGs (MAGs 2, 6, 8, 9, 10, 11, 13, 14, 15, 16, 21, 26, 27, and 29), and only MAG 17 was fermentation dominated. Notably, no MAG was dominated by hydrocarbon degradation as a top category. However, it was a connecting node across all 26 MAGs.

*Pseudomonadales* MAGs (n = 12) revealed consistent connectivity across three metabolic categories, fermentation, central carbon metabolism, and hydrocarbon degradation, with all MAGs exhibiting three edges, indicating a uniform and fully connected network topology across this order (Figure 7b, Table S6). Total edge weights ranged from 0.85 to 2.43, reflecting variation in the strength of metabolic co-expression among individual MAGs. Fermentation was the dominant category in 7 MAGs (MAGs 3, 8, 15, 16, 17, 18, and 19), while central carbon metabolism led in the remaining 5 MAGs (MAGs 1, 2, 5, 6, and 21). Hydrocarbon degradation was not dominant as a top category for any MAG, however, its presence as a connected node across all 12 MAGs indicates that hydrocarbon degradation is universally co-expressed alongside fermentation and central carbon metabolism.

Metatranscriptome analysis revealed distinct co-metabolism patterns that varied by bay, season, and salinity (Figure S4, Table S6). *Burkholderiales* MAGs (n= 26) exhibited bay, salinity and season dependent metabolic coupling. Under summer low salinity conditions in both bays, hydrocarbon degradation, C1 metabolism, and central carbon metabolism genes were co-expressed expressed (e.g., MAG 4). In contrast, during spring high and mid salinity conditions across both locations, hydrocarbon degradation co-occurred with central carbon metabolism (e.g., MAG 13). Additional MAGs showed hydrocarbon degradation and central carbon metabolism co-expression during summer low salinity conditions in both bays (MAGs 8 and 14), as well as fall low salinity conditions in Delaware Bay and spring low salinity conditions in Chesapeake Bay (e.g., MAG 16). *Pseudomonadales* (n = 12) displayed bay, salinity and season structured metabolic integration. Hydrocarbon degradation, fermentation, and central carbon metabolism genes were co-expressed during spring mid and high salinity conditions in Chesapeake Bay, with similar patterns observed in summer mid salinity conditions (e.g., MAG 1). This tripartite co-expression also occurred in summer mid to high salinity conditions in Chesapeake Bay and fall high salinity in Delaware Bay, with summer high salinity Delaware Bay samples showing comparable patterns (e.g, MAG 2). In both bays, hydrocarbon degradation co-occurred with central carbon metabolism and fermentation during spring mid to high salinity conditions (MAGs 15 and 18). Across all active MAGs in both *Burkholderiales* and *Pseudomonadales*, genes encoding key metabolic functions including respiration were consistently co-expressed with primary catabolic pathways.

## Discussion

Our analysis demonstrated that the two estuarine systems studied harbor taxonomically diverse microbial assemblages with hydrocarbon degradation potential, whose composition, metabolic potential, FRed and transcriptional activity are structured by environmental gradients. By linking genome resolved metabolic potential with community wide gene transcription patterns, these findings address critical gaps in understanding how environmental conditions shape the distribution and activity of hydrocarbon degrading taxa in estuarine waters. Environmental gradients, particularly salinity, structured both community composition and pathway specific gene expression. FRed varied substantially across pathways, with catechol degradation showing high redundancy distributed across many orders while specialized pathways like naphthalene and cymene degradation remain taxonomically restricted and potentially vulnerable. Co-expression analysis revealed complementary metabolic strategies of condition dependent niche partitioning and integrated metabolism, with both strategies operating at multiple scales to support ecosystem resilience. This integrated genomic framework provides a mechanistic basis for predicting microbial community responses to changing hydrocarbon loads, seasonal environmental shifts, and anthropogenic disturbance in coastal ecosystems, with direct implications for bioremediation strategies and coastal ecosystem management under increasingly variable conditions.

### Diverse Hydrocarbon Degrading Community Across Estuaries

Estuarine microbial communities in the Chesapeake and Delaware Bays harbor a diverse array of aerobic and anaerobic hydrocarbon degrading taxa spanning 41 orders, highlighting the functional versatility of these communities. The dominant hydrocarbon degrading orders of *Burkholderiales*, *Pseudomonadales*, *Flavobacteriales*, *Acidimicrobiales*, *Rhodobacterales*, and *Nanopelagicales* exhibited broad metabolic potential, encoding enzymes for both aliphatic and aromatic hydrocarbon degradation. These orders are frequently detected in petroleum or PAH impacted environments, consistent with observations in various marine and freshwater systems (8, 44–46).

### Environmental Gradients Structure Community Composition and Transcriptional Activity

Environmental gradients strongly influenced both community composition and hydrocarbon degradation potential, with salinity as the primary driver, followed by temperature and nutrients. These trends are consistent with the established microbial compositions in estuaries, driven mainly by salinity (33). Core taxa such as *Burkholderiales* and *Pseudomonadales* were present across low and mid/high salinity regimes, respectively, mainly in the spring but there were many more orders present in the summer. Metatranscriptomic analysis revealed that *Pseudomonadales* dominated transcriptional activity in spring, particularly in the Chesapeake Bay, while *Burkholderiales*, *Flavobacteriales*, and *Rhodobacterales* MAGs were most active during summer and fall under medium and high salinity conditions. These seasonal patterns are consistent with known temperature-dependent constraints on hydrocarbon degradation, where warmer periods (20–30 °C) promote enhanced enzyme activity and faster PAH turnover, as documented in temperate and subtropical systems (45, 47). This combination of core and dynamic taxa likely supports both stability and adaptability in hydrocarbon degradation under fluctuating estuarine conditions, consistent with patterns observed in other hydrocarbon impacted coastal systems (45, 48, 49).

### Hydrocarbon Sources and Physicochemical Conditions Drive Pathway Specific Distributions

The distribution of hydrocarbon degradation genes reflected similar environmental trends at the community level. Naphthalene, xylene, biphenyl and catechol degradation genes showed different correlations with salinity, nutrients and temperature, highlighting pathways that are specifically responsive to environmental change. Salinity was positively correlated with aromatic hydrocarbon degradation genes, while availability of nutrients and temperature was correlated with subsets of genes involved in the alkane and xylene pathways, in line with findings in other anthropogenic affected estuaries (50).

Specifically, catechol meta-cleavage pathway genes exhibited significant positive correlations with salinity and strong negative correlations with temperature, silicate, nitrate, and phosphate concentrations, indicating these genes are most abundant under higher salinity, lower temperature and lower-nutrient conditions. In contrast, naphthalene degradation genes showed strong negative correlations with salinity and strong positive correlations with nitrate and phosphate, indicating these pathways are most abundant in nutrient-rich, low-salinity upper estuarine waters. Similarly, xylene degradation genes exhibited negative correlations with salinity and positive correlations with nitrate and phosphate. These pathway specific nutrient associations demonstrate that nutrient availability modulates degradation efficiency in a context-dependent manner, with both nutrient limitation and nutrient excess capable of suppressing specific PAH degradation pathways, consistent with biostimulation studies in diverse systems (51–53).

The pronounced differences in hydrocarbon degradation gene expression between Delaware and Chesapeake Bays align with distinct hydrocarbon sources and physicochemical regimes. Delaware Bay showed consistently higher expression of both the alkane and aromatic degradation pathways, in particular the genes for ortho- and meta-cleavage of catechol, benzene and xylene, with the highest transcriptional activation in the low salinity spring samples. This is likely due to intensive anthropogenic PAH loading of the bay from highly urbanized and industrial corridors, which are delivered by storm water runoff and river transport during peak discharge seasons (54).. The high turbidity in Delaware Bay, where resuspended inorganic sediments dominate, further biases the system toward allochthonous, combustion-derived aromatic hydrocarbons rather than phytoplankton derived aromatics (55). The Chesapeake Bay, on the other hand, showed a lower overall abundance, which is consistent with its lower turbidity and a greater contribution from natural marine and aquatic aromatic compounds (56, 57). Moderate spring activation followed by summer decline is consistent with thermal stratification and hypoxia developing, which may be a factor in the inhibition of aerobic degradation (58, 59). In both bays, the low salinity samples showed the highest abundance and expression of genes, which reflects the spatial concentration of pyrogenic PAHs in upper estuarine waters during high freshwater discharge. The seasonal peaks in spring correspond to high water runoff, while the sustained activity in summer and fall reflects the continuous input of pollutants combined with warmer temperatures increasing microbial metabolism [60].

Aerobic aromatic degradation genes were consistently the most abundant in both estuaries, whereas aerobic alkane degradation genes represented a substantial secondary component. Anaerobic hydrocarbon genes were relatively rare and limited to specific seasons and salinity conditions. The dominance of aerobic aromatic degradation genes is consistent with evidence that surface estuarine environments tend to support oxygen-rich conditions that facilitate dioxygenase and monooxygenase mediated ring oxidation, as observed across diverse marine systems (59, 60). . By contrast, the relative scarcity of anaerobic hydrocarbon genes corroborates the report that anaerobic PAH degradation requires specific redox conditions and tends to occur primarily in deeper sediments or aquifers.

### FRed and Co-metabolism Support Ecosystem Resilience

The distribution of hydrocarbon degradation genes across taxonomic lineages revealed substantial variation in FRed with important implications for ecosystem stability. Catechol degradation pathways exhibited the highest redundancy, with meta-cleavage genes found in 67% of MAGs and ortho-cleavage genes in 41% of MAGs, suggesting that core aromatic degradation capacity is deeply embedded across estuarine bacterial communities and likely buffers against environmental perturbations (61). Metatranscriptomic analysis revealed that potential redundancy translated into active FRed under favorable conditions (62, 63). In Delaware Bay spring low salinity samples, MAGs from multiple orders simultaneously expressed catechol degradation genes, demonstrating realized FRed wherein phylogenetically diverse lineages actively contribute to the same ecosystem function (40). However, environmental stresses constrained this redundancy, high salinity conditions suppressed expression across multiple orders simultaneously, suggesting that environmental filters can create functional bottlenecks despite persistent genetic potential (39). In contrast, specialized pathways exhibited minimal redundancy suggesting vulnerability where loss of specific taxa could compromise ecosystem level degradation capacity.

Beyond taxonomic and FRed, co-expression analysis revealed complementary metabolic strategies that enhance functional resilience through distinct mechanisms. Both *Burkholderiales* and *Pseudomonadales* exhibited bay, salinity, and season dependent metabolic coupling wherein hydrocarbon degradation genes were consistently co-expressed with central carbon metabolism genes and key metabolic genes across environmental gradients. However, they differed fundamentally in their metabolic integration strategies. *Burkholderiales* displayed condition specific pathway activation wherein C1 metabolism genes were co-expressed with hydrocarbon degradation and central carbon metabolism genes only under summer low salinity conditions when osmotic stress is reduced, but not during spring high and mid salinity conditions. This heterogeneous, condition dependent co-expression profile suggests niche partitioning wherein different MAGs within *Burkholderiales* occupy distinct metabolic niches (64)., providing finer scale redundancy where loss of one MAG may be compensated by another with overlapping but distinct metabolic capabilities. In contrast, *Pseudomonadales* displayed metabolic integration wherein hydrocarbon degradation genes were uniformly co-expressed with both fermentation and central carbon metabolism genes across spring mid and high salinity conditions in Chesapeake Bay, summer mid to high salinity conditions in both bays, and fall high salinity in Delaware Bay. Although fermentation is associated with anaerobic conditions, its co-expression in *Pseudomonadales* surface water MAGs is consistent with the well-documented capacity of *Pseudomonas* and related taxa to employ fermentative overflow metabolism aerobically, observed when carbon flux through glycolysis exceeds the respiratory capacity of the cell, resulting in the excretion of fermentation end products such as acetate and formate even under oxic conditions (65). In addition, estuarine surface waters do not have uniform oxygen environments, and micro-organisms in organic matter aggregates, biofilms and during periods of high biological oxygen demand associated with seasonal productivity pulses (66).create conditions favorable to the development of fermentation pathways and the production of redox and ATP. The co-expression of fermentation with hydrocarbon degradation is therefore likely to reflect a facultative metabolic strategy whereby *Pseudomonadales* may be able to dynamically switch between full- and mixed-phase fermentation of carbon in response to micro-scale variations in oxygen and substrate in surface waters. This constitutive tripartite co-expression indicates metabolic versatility, conferring resilience through metabolic flexibility where individual populations can shift between carbon sources as availability changes, maintaining ecosystem function without requiring compensatory responses from other taxa (67).

As mentioned in the previous paragraph, genes encoding key metabolic functions including respiration were consistently co-expressed with primary catabolic pathways. These complementary strategies of *Burkholderiales* within-lineage diversity and *Pseudomonadales* metabolic integration, jointly contribute to robust hydrocarbon degradation capacity, aligning with portfolio effect theory wherein diversity in functional strategies enhances community level stability (68). The persistence of multiple active lineages across environments, coupled with diverse co-metabolic strategies, demonstrates that FRed operates at multiple taxonomic scales, realized expression under favorable conditions, and coordinated pathway activation within individual MAGs. In hydrocarbon rich ecosystems, this multi-scale redundancy enables overlapping taxa to maintain degradation processes even when environmental perturbations alter species composition, consistent with observations following the Deepwater Horizon spill (49). This redundancy likely contributes to the capacity of estuarine microbial communities to respond rapidly to episodic hydrocarbon inputs while maintaining baseline degradation activity under normal conditions.

### Conclusion and Broader Implications

Our findings indicate that the hydrocarbon degrading microbial communities in Chesapeake and Delaware Bay are taxonomically diverse, environmentally structured and functionally resistant, with salinity, temperature and seasonality being the major drivers of both the composition of the community and the expression of the in-situ degradation genes. These findings advance the understanding of hydrocarbon degradation in estuarine ecosystems, which are major reservoirs and transition zones for PAH and other hydrophobic organic contaminants, and provide a mechanistic framework with a genome resolution to predict degradation potential under changing environmental conditions, including climatic changes in salinity and temperature, nutrient loading and episodic hydrocarbon inputs. These results are also directly relevant to bioremediation by identifying key adaptable taxa, co-metabolic strategies for specific conditions and environmental windows that support optimal degradation. Such knowledge can inform nature-based strategies for mitigating hydrocarbon contamination in coastal systems, as demonstrated in previous biostimulation experiments (47, 69). Collectively, this work highlights that hydrocarbon degradation resilience is not uniformly distributed across pathways and highlights the need for genome-based, pathway-specific approaches to accurately predict and manage microbial degradation capacity under increasing anthropogenic and climate pressures.

## Methodology

### Site Description and Sample Collection

Surface water samples were collected from longitudinal transects of the Delaware Bay (spring, summer, fall 2014) and Chesapeake Bay (spring, summer 2015), spanning a salinity gradient of 0.08–30.52 and depths of 1.2–4.47 m. Samples were sequentially filtered through 0.8 μm and 0.22 μm pore-size filters to obtain two microbial size fractions. Environmental metadata (temperature, salinity, depth) and GPS coordinates were recorded at each station. Full details of sampling and site characterization are described elsewhere (38, 70) and in supplementary information.

### Metagenomic and Metatranscriptomic Sequencing

DNA and RNA were co-extracted from filters, and library preparation, sequencing, and data processing were performed at the Joint Genome Institute as previously described (38, 70). In total, 36 metagenomic and 58 metatranscriptomic libraries were sequenced. Reads were quality-trimmed, assembled, and binned following established workflows, resulting in 360 dereplicated metagenome-assembled genomes (MAGs; ≥95% ANI) (41, 70). Additional details of sequencing and assembly procedures are provided elsewhere (38, 70).

### Functional Annotation of MAGs

Metagenome-assembled genomes (MAGs) were annotated using DRAM v1.4.6 (Distilled and Refined Annotation of Metabolism) (71) to provide genome resolved metabolic reconstruction and context for pathway completeness within individual populations. DIAMOND v2.0.14 (72) was used for read level homology searches to quantify marker gene abundance and transcription across samples with high sensitivity. Based on this screening, MAGs containing at least one hydrocarbon-degradation marker were identified and selected for downstream analyses. For more details see supplementary information.

### Abundance and Functional Profiling of Hydrocarbon-Degrading Bacteria

MAG abundances in metagenomes and metatranscriptomes were calculated by mapping reads using Bowtie2 v2.2.5.4 (73) and converting SAM files to BAM format with SAMtools v1.21.1 (74). Feature counts were generated using Subread v2.0.8 (75). Gene-level metagenomic counts were normalized as Reads Per Kilobase per Genome Equivalent (RPKG), accounting for gene length and genome equivalents (76), while metatranscriptomic counts were normalized to Transcripts Per Million (TPM) by scaling to taxonomy by sum of genome lengths (77). MAG-level abundances were normalized as Counts Per Million (CPM) using CoverM v0.6.1 (78), calculated as the proportion of reads mapping to each dereplicated MAG relative to total Kaiju-classified bacterial and archaeal reads across Chesapeake Bay and Delaware Bay metagenomes (41).

### FRed, Co-metabolism and statistical analysis

All statistical analyses were performed in R v4.5.0 (79). Correlations between hydrocarbon degradation genes, environmental parameters, and community composition were evaluated using Spearman’s rank correlation with p-adjusted values. Ordination analyses were conducted with distance base redundancy analysis (dbRDA) using the vegan package to assess relationships between microbial communities and environmental gradients (80).

To distinguish between potential and realized Fred, metagenomic and metatranscriptomic feature count tables were compared by quantifying pathway-level hits for each taxonomic order. Orders contributing hits to the same hydrocarbon degradation pathways across multiple taxa were considered functionally redundant, with metagenomics capturing encoded capacity and metatranscriptomics revealing which of those functions were actively expressed *in situ*.

Metabolic co-activity analyses focused on the prominant hydrocarbon-degrading taxa, *Burkholderiales* (n = 26) and *Pseudomonadales* (n = 12), across environmental conditions with at least one gene from five metabolic categories (hydrocarbon degradation, C1 metabolism, complex carbon degradation, fermentation, and central carbon metabolism). Co-metabolism was quantified using log₂-transformed sums of gene expression values from five metabolic categories per MAG. The Variance-stabilized expression values were used to make the network analysis in R using dplyr. Co-energy utilization was assessed by screening the selected MAGs against custom HMM-based databases targeting marker genes for respiration, sulfur cycling, nitrogen cycling, carbon fixation, and phototrophy (41). Metabolic potential was inferred from marker gene presence in metagenomes, and transcriptional activity was quantified as TPM from feature-counted GFF-mapped metatranscriptomic reads.

## Acknowledgements

We gratefully acknowledge the crew of the R/V Hugh R. Sharp for their contributions to sample collection, and David Kirchman, Matt Cottrell, and Liying Yu for their invaluable technical expertise and sampling assistance. Computational analyses were performed on the Palmetto Cluster at Clemson University, supported by the National Science Foundation (MRI# 2024205, MRI# 1725573, and CRI# 2010270). We also acknowledge the Clemson University Genomics and Bioinformatics Facility for bioinformatics support, funded in part through Institutional Development Awards (IDeA) from the National Institute of General Medical Sciences of the National Institutes of Health (P20GM146584 and P20GM139769).

B.J.C. conceptualized the study, secured funding, and directed field sampling efforts. M.O. performed MAG binning. J.J. contributed to study design and analyses. D.J. conducted all data analyses and prepared the original manuscript draft under the supervision of B.J.C. All authors critically reviewed and approved the final manuscript.

## Conflict of Interest

No conflicts of interest

## Funding

Funding for the research cruises, metagenomic and metatranscriptomic sequencing, and data processing was provided by the National Science Foundation (OCE-082546 and EF-2025541) and the DOE/JGI (CSP-1621), all awarded to B.J.C.

## Data Availability

The metagenomes, metatranscriptomes, and MAGs are available in NCBI under the umbrella project PRJNA432171.

All the detailed codes and calculations are available on our GitHub page: https://github.com/Campbelllab-bioinfo under Environmental-Gradients-Shape-the-Hydrocarbon-Degrading-Microbiome-in-Two-Mid-Atlantic-Bays.

## Supplementary Information

**Supplementary Materials and Methods.** A more detailed materials and method section of the main text.

**Figure S1**. Abundance of hydrocarbon degradation genes within metagenome across the samples, quantified as log_10_ CPM.

*Abbreviations:* CP = Chesapeake Bay; DE = Delaware Bay; Spr, Sum, Fall = Spring, Summer, Fall; G08 = >0.8 μm; L08 = <0.8 μm; CPM = counts per million. Full gene names, enzyme functions, and pathway details are provided in Supplementary Table 1.

**Figure S2.** Phylogenomic tree of 181 dereplicated MAGs constructed using 43 conserved single-copy marker genes identified by CheckM. The background color of each clade indicates the taxonomic order to which each MAG belongs as assigned by GTDB-Tk. Bar charts at tree tips indicate the CPM-normalized metagenomic abundance of each MAG across Chesapeake Bay and Delaware Bay metagenomes.

**Figure S3.** Metatranscriptome-derived hydrocarbon degradation gene expression, quantified by log_2_ TPM-normalized transcript abundance values.

*Abbreviations:* CP = Chesapeake Bay; DE = Delaware Bay; Spr, Sum, Fall = Spring, Summer, Fall; TPM = transcripts per million. Full gene names, enzyme functions, and pathway details are provided in Supplementary Table 1.

**Figure S4.** Co-expression patterns of metabolic genes across *Burkholderiales* and *Pseudomonadales* MAGs in Chesapeake Bay and Delaware Bay metatranscriptome. Heatmaps display normalized log2 expression of genes grouped by key metabolic process.

**Table S1.** CPM-normalized metagenomic abundances of hydrocarbon degradation genes identified by DIAMOND search across all Chesapeake Bay and Delaware Bay metagenome samples.

**Table S2.** KEGG Orthology (KO) identifiers, gene names, and associated hydrocarbon degradation pathways for genes used in metagenomic and metatranscriptomic analyses.

**Table S3.** Number of MAGs per taxonomic order encoding genes involved in each hydrocarbon degradation pathway across metagenomic samples. Values indicate the total number of MAGs per order, with pathway totals summarized by total orders and total MAGs.

**Table S4.** Summary of distance-based redundancy analysis (dbRDA) results examining the influence of environmental variables on the abundances of hydrocarbon degradation-related MAGs across Chesapeake Bay and Delaware Bay metagenomes. Degrees of freedom, sum of squares, F-values, and significance levels are reported for the overall model and individual parameter.

**Table S5.** Number of MAGs per taxonomic order encoding genes in each hydrocarbon degradation pathway across metatranscriptomic samples, summarized by total orders and total MAGs per pathway.

**Table S6.** Presence/absence matrix of key metabolic marker genes across all MAGs, gene ID reference list, and summary of co-expression network analysis of *Burkholderiales* and *Pseudomonadales* MAGs.

